# Dietary fatty acids as determinants of cuticular wax profiles in *Tribolium castaneum*

**DOI:** 10.1101/2025.07.06.662788

**Authors:** Subhadeep Das, Avishek Dolai, Oishika Chatterjee, Riya Saha, Pritom Das, Sourav Manna

**Author notes:** Correspondence - Sourav Manna, Department of Life Sciences, Presidency University, Kolkata, 700073, India.

## Abstract

Cuticular waxes are complex mixtures of hydrocarbons and other lipophilic compounds that form an outer protective layer over the insect cuticle playing a role in ecological interactions as well as physiological processes. While CHC profiles are largely genetically determined, several studies have linked diet to cuticular hydrocarbons and limited research indicates a potential influence of dietary fatty acids. Considering this, the current work investigates the effect of nine cereal flours, i.e., wheat, rice, maize, barley, oat, ragi, sorghum, soybean and gram, each with a distinct fatty acid composition, on the cuticular wax profile of *Tribolium castaneum* (Coleoptera: Tenebrionidae), a globally significant stored grain insect. Fatty acid compositions of the nine flours and the cuticular waxes from the beetle populations reared on each diet were characterized using gas chromatography mass spectrometry (GC-MS). In all the beetle populations, the most abundant compound was 1-pentadecene, followed by cis-9-tetradecen-1-ol and 1-heptadecene, irrespective of the diet. While no qualitative differences in cuticular wax composition were found, significant quantitative variation in cuticular wax profile was observed across dietary groups. Similarity percentage (SIMPER) analysis revealed a cumulative Bray– Curtis dissimilarity of 47.72 among the flour samples based on their fatty acid profiles, and a dissimilarity of 18.51 among *T. castaneum* populations reared on these flours based on their cuticular wax composition. Non-metric multidimensional scaling (NMDS) further indicated substantial variation in the cuticular wax profiles of beetle populations raised on different dietary sources. Pearson’s correlation analysis between the concentrations of individual cuticular wax components and the fatty acid content of the respective diets demonstrated strong correlations between specific cuticular compounds and dietary fatty acids. Notably, levels of 1-pentadecene in the beetle cuticle were strongly associated with the concentrations of hexadecanoic and octadecanoic acids in the flours. These results support the hypothesis that, alongside endogenous conserved biosynthetic pathways, dietary fatty acid composition contributes to the modulation of insect cuticular chemistry. This dietary influence may have important implications for understanding insect ecological plasticity and diet-mediated changes in surface chemistry.

## Introduction

Cuticular waxes, present on the cuticular surfaces of insects play multidimensional role in physiology and behaviour of insects. Multiple pieces of literature highlighted on the role of cuticular substances to prevent desiccation in higher temperature (Blomquist et al., 2018; Gibbs, 2007). In addition, these substances are also being increasingly recognized for their role in intra and interspecific chemical communication (Blomquist et al., 2018; Ferveur, 2005). Cuticular waxes comprise a range of classes of molecules such as hydrocarbons, long chain alcohols, long chain aldehydes and long chain acids (Buckner et al., 1999), of which hydrocarbons, and particularly cuticular hydrocarbons, are the most abundant (Gołębiowski and Stepnowski, 2022). The composition and diversity of cuticular lipids, particularly hydrocarbons, expressed on the cuticle of a given taxon are typically under genetic control (Ferveur and Jallon, 1996) and thus, often can be used as taxonomic identification of a species (Moore et al., 2021). However, their quantitative variation may be influenced by various biotic factors, such as sex, age, degree of parasitism, and abiotic factors such as temperature (Blomquist and Ginzel, 2021; Walsh et al., 2020).

Although cuticular waxes, particularly cuticular hydrocarbons, exhibit considerable diversity across insect taxa, their fundamental biosynthetic pathway remains conserved. Insects manufacture the majority of compounds in their cuticular wax by themselves (Nelson and Buckner, 1995; Blomquist and Bagnères, 2010), with much lower amounts of dietary hydrocarbons being directly absorbed into their epicuticle (Blomquist and Jackson, 1973a). These cuticular hydrocarbons are believed to be synthesized via fatty acid metabolism, involving key enzymes such as fatty acid synthase, fatty acyl-CoA elongase, and fatty acyl-CoA reductase, which are predominantly active in oenocyte cells or fat bodies (Blomquist et al., 2018). The synthesized hydrocarbons are subsequently shuttled to the cuticular surface through the haemolymph by high-density lipoprotein lipophorin (Blomquist and Bagnères 2010). The fatty acids that are involved in the biosynthesis of hydrocarbons are mostly produced via de novo fatty acid biosynthesis pathway by the condensation of malonyl-CoA units with acetyl-CoA that produces fatty acyl CoA, which eventually via enzymatic modification transform into structurally diverse CHCs (Blomquist and Bagnères, 2010; Howard and Blomquist, 2005).

Even though expression of CHCs are genetically determined, evidences suggest that diet can significantly influence the relative ratios of hydrocarbons in the cuticular surface in multiple insect species, such as in *Linepithema humile* (Hymenoptera) (Buczkowski and Silverman, 2005; Liang and Silverman, 2000a; van Wilgenburg et al., 2022a), *Manduca sexta* (Lepidoptera) (Espelie and Bernays, 1989), *Melanoplus sanguinipes* (Orthoptera) (Blomquist and Jackson, 1973b), *Phaedon cochleariae* (Coleoptera) (Geiselhardt et al., 2012). There are evidences that suggest that certain species incorporate the dietary hydrocarbons into the cuticular lipid, such as in *M. sanguinipes* (Blomquist and Jackson, 1973b). On the other hand few literatures reveal that diet can alter the gut microflora which can indirectly influence the cuticular hydrocarbon profile, such as in *Drosophila melanogaster* (Diptera) (Sharon et al., 2010). Research has been exploring the function of dietary macromolecules in defining cuticular chemical composition, and it was found that dietary carbohydrates and amino acids might not contribute in the biosynthesis and subsequent expression of hydrocarbons in the cuticular surface of insects. Rather, it is believed that dietary fatty acids play an important role in determining the relative abundances of these hydrocarbons. In *P. cochleariae*, it has been reported that the consumption of different fatty acid blends leads to quantitative changes in the beetle’s straight-chain and methyl-branched CHCs (Otte et al., 2015). Pennanec’h et al. (1997) also hinted towards similar phenomenon in *D. melanogaster*. However, there is still limited literature available to enhance our understanding of whether and how fatty acids influence the determination of an insect’s cuticular hydrocarbon profile.

Red flour beetle of *Tribolium castaneum* (Coleoptera: Tenebrionidae) is a well-known pest of stored grain products and thrives on various cereal-based flours, causing significant deterioration in food quality by degrading essential macronutrients (Das et al., 2024). In addition to its economic importance, this species is also widely recognized as a model organism in entomological studies, particularly in both fundamental and applied research (Campbell et al., 2022). Therefore, *T. castaneum* was selected for the present study to investigate whether dietary fatty acids play a significant role in determining the composition and abundance of cuticular surface compounds. To address this, adults were reared for three consecutive generations on nine distinct cereal flours, each assumed to possess a unique fatty acid profile. The composition of cuticular wax of these beetles was subsequently analyzed to evaluate potential associations between the fatty acid composition of the diet and the chemical profile expressed on the beetles’ cuticle. This study aimed to answer the research question that if there is an association between the dietary fatty acid compositions of flours and the cuticular chemical composition of *T. castaneum*.

## Materials and methods

### Insect Rearing

Whole grain flours, including wheat, rice, chickpea, corn, finger millet, sorghum, pearl millet, Job’s tears, and oat, were purchased from a local mill. To eliminate any potential contamination by *Tribolium castaneum* (eggs, larvae, or pupae) or other insect pests, all flours were sterilized in a hot air oven at 60°C for 30 minutes (Das et al., 2024). Fifteen grams of each flour were measured and placed into 100 mL round plastic containers. In each container, thirty 7-day old adult *T. castaneum* (15 males and 15 females) were introduced. The cultures were maintained under controlled conditions: a 12:12 hour light–dark photoperiod at 25 ± 1 °C and 65 ± 3% relative humidity. To minimize the effects of deteriorating nutritional and fatty acid quality, which could influence experimental outcomes, all individuals were carefully transferred to respective fresh flour every seven days. This weekly transfer also served to prevent population growth, as an increase in population size could alter the composition of cuticular compounds. The experiment was conducted for three months. At the end of the three-month period, five individuals were randomly selected from each container and pooled as a single biological replicate. For each of the nine flour types tested, three such biological replicates were prepared.

### Extraction of lipids and Fatty acid Analysis

Bligh and Dyer’s method were used to extract lipids from freshly procured nine flours (Bligh and Dyer, 1959). 200 mg of each flour was taken in a separating funnel and Methanol, Chloroform and the water were combined to make a mixture in the following proportion-2:1:1 (by vol). Finally, distilled water and chloroform were each added at one-third of the total sample/mixture volume to facilitate phase separation. The separating funnel was shaken time to time for phase separation and stored in the refrigerator in an undisturbed state for 72 hours. The lipid rich lower chloroform phase was taken out, collected in a screw-capped septum vial, and then anhydrous sodium sulphate was added to it before being refrigeration to 4°C for further analysis. Fatty acids (FAs) present in the lipid fractions (chloroform phase) were transesterified to form fatty acid methyl esters (FAME) (Christie, 1993). For total lipid, 1 mL of purified chloroform phase was added to a 5mL solvent mixture of methanol, benzene, and conc. H_2_SO_4_ (4.3:0.5:0.6, by vol), and the mixture was incubated at 90°C for 8 h and left overnight at room temperature. Before esterification, 1 µL of internal standard, n-Decanoic acid (1mg/ml) was mixed in chloroform phase. Following esterification, 4–5 mL of hot distilled water was added to the reaction mixture, and vigorous mixing was performed using a Pasteur pipette in a test tube. Saponifiable containing FAMEs were eluted with n-hexane (E. Merck, India, HPLC grade) and finally dried over anhydrous sodium sulphate. The n-hexane was reduced in volume under a nitrogen flow and then analysed using Gas Chromatography-Mass Spectrometry (GC-MS).

### Analysis of cuticular waxes/hydrocarbons

To extract cuticular wax compounds, five adult beetles were immersed in 200 µL of n-hexane for 5 minutes. The resulting extracts were concentrated to 50 µL under a mild nitrogen stream. Subsequently, 1 µL of nonane (100µg/mL) was added to the extract as an internal standard. The final mixture was then injected into the GC-MS for analysis.

### GC-MS parameters for chemical analysis

Thermo Fisher Trace 1300 (Thermo Scientific, Milan, Italy) GC-MS was conducted using a triple-quadrupole mass spectrometer (Thermo MS-TSQ 9000) fitted with a wall-coated open tubular column (WCOT), TG-5 MS (30 m × 0.25 mm × 0.25 μm), which was connected to a 10 m Dura guard capillary column. GC-MS analysis was conducted with the front inlet temperature set at 250°C. Helium, with a purity of better than 99.99%, acted as the carrier gas at a flow rate of 1 mL/min.

### GC-programme for analysis of FAME

The initial oven temperature was set at 125°C with 2 min hold time, followed by a ramp increase at a rate of 5°C/min until reaching 250°C. A final hold at 250°C was maintained for 8 minutes. Total run time was 35 minutes.

### GC-programme for cuticular wax

The initial oven temperature was set at 50°C with 1 min hold time, followed by a ramp increase at a rate of 6°C/min until reaching 300°C. A final hold at 300°C was maintained for 8 minutes. Total run time was 51 minutes.

For all of the above programme, the transfer line temperature of the mass spectrometer was maintained at a constant 250°C, and the ion source temperature was set at 220°C. The mass spectrometer operated in electron ionization (EI) mode with an electron energy of 70 eV.

The data were processed using “Chromeleon Chromatography Data System (CDS) software” from Thermo Fisher Scientific, and identification of FAME and n-alkanes present in cuticular wax was done by comparing with m/z mass spectrum of the individual compounds with NIST 2020 mass spectral library, as well as comparing the retention time of authentic analytic standards procured from Supelco (LB80556 and LB77207, USA) and Sigma Aldrich (Lot no: 40147-U) respectively. For components of the cuticular wax that did not belong to the n-alkane category, identification was carried out not only by referencing the NIST spectral library but also by calculating their linear retention indices (LRI). The LRI for any specific compound was estimated as:

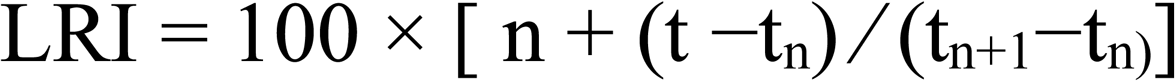

[Where n corresponds to the carbon number of the n-alkane that immediately precedes the target compound. t is the retention time of the compound, whereas t_n_ and t_n+1_, represent the retention times of the n-alkanes that directly precede and follow the VOC, respectively. These retention times are determined under identical experimental conditions, using the same temperature gradient and chromatographic column setup.].

For GC-MS data interpretation, total ion current (TIC) area of each identified compound was normalized with the area of internal standards, decanoic acid for FAME analysis and n-nonane for cuticular compound analysis. The final concentrations of the compounds are represented in the manuscript as µg/g flour (for FAMEs) and µg/insect (for cuticular compounds) respectively.

### Statistical analysis and graphical representation

All data obtained from the aforementioned analyses were first subjected to the Shapiro–Wilk test to assess the normality of distribution. Given the non-normal nature of the data, the Kruskal–Wallis test was employed to determine statistically significant differences between the tested groups based on specific variables. Post hoc comparisons were conducted using Dunn’s Sidak test. To assess the relationship between the concentration of specific fatty acids in flour samples and the concentration of corresponding cuticular compounds in *T. castaneum* wax, Pearson’s correlation analysis was performed. A p-value of <0.05 was considered the threshold for statistical significance in all tests.

For multivariate analyses, Similarity Percentage (SIMPER) analysis was conducted using Bray–Curtis dissimilarity to quantify the overall dissimilarity among flour samples (based on fatty acid composition) and *T. castaneum* populations (based on cuticular wax profiles). SIMPER also identified the key variables (e.g., specific fatty acids or cuticular compounds) that contributed most to the observed differences between predefined groups. Non-metric Multidimensional Scaling (NMDS) was performed to visualize the Bray–Curtis dissimilarity between sample groups, both for flour samples (based on fatty acid profiles) and *T. castaneum* populations (based on cuticular wax composition). For factor analysis, the fatty acid composition of flour samples was treated as an environmental variable to evaluate whether these compounds could explain the clustering pattern of *T. castaneum* populations based on their cuticular wax profiles.

All statistical analyses and graphical representations were carried out using OriginPro 2024b or PAST 5, depending on analytical requirements.

## Results

### Fatty acid composition of dietary source

The analysis of fatty acids from the investigated flour samples revealed the presence of fatty acids ranging from tetradecanoic acid (C14) to tetracosanoic acid (C24), with considerable variation observed in the fatty acid profiles among the different flours (Figure 1a; Table 1). In proportional terms, hexadecanoic acid was one of the most dominant fatty acids in most samples—especially wheat, rice, corn, finger millet, pearl millet, Job’s tears, and oat flour. However, its absolute concentration varied considerably among these flours. For instance, rice flour contained 33–52 µg/g and wheat flour 54–78 µg/g of hexadecanoic acid, while oat flour exhibited an exceptionally high concentration ranging from 404–428 µg/g (Figure 2a; Table 1). Unsaturated fatty acids also constituted a substantial proportion of the total fatty acids in many of the flours, with the exception of rice and wheat. In particular, chickpea, corn, sorghum, and pearl millet flours showed a higher ratio of unsaturated to saturated fatty acids. Among the unsaturated fatty acids, 9-octadecenoic acid, 11-octadecenoic acid, and 9,12-octadecadienoic acid were the most abundant across all analyzed samples. In addition to these, octadecanoic acid and eicosanoic acid were detected in considerable quantities in most samples (Figure 2a).

**FIGURE 1.**
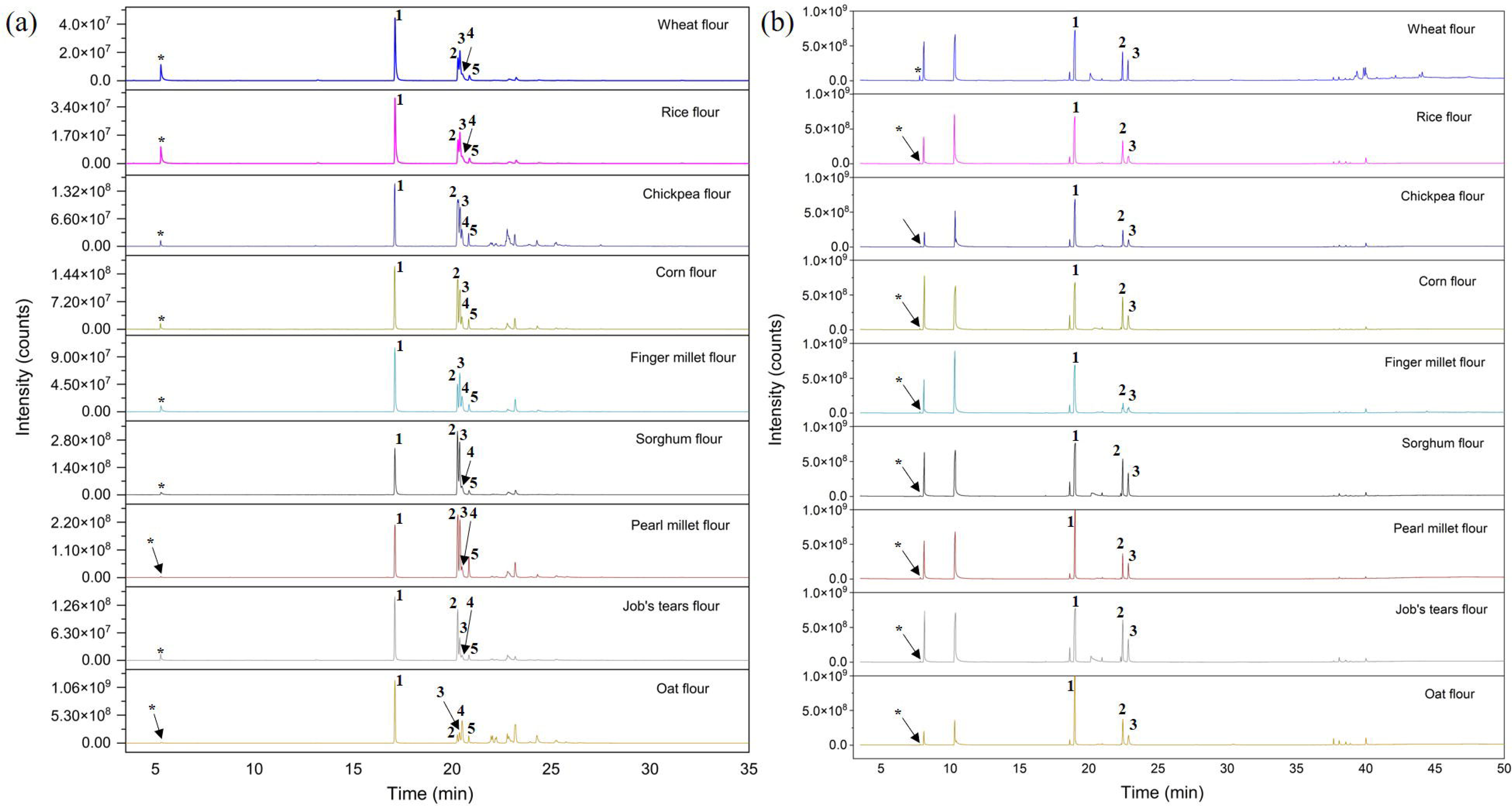
GC chromatogram of the fatty acid profiles of the nine flour types (a); the asterisk (*) denotes the internal standard. Peaks labelled 1, 2, 3, 4, and 5 correspond to five major fatty acids: 1 – Hexadecanoic acid, 2 – Linoleic acid, 3 – Oleic acid, 4 – Elaidic acid, and 5 – Octadecanoic acid. GC chromatogram showing the cuticular wax profile of *Tribolium castaneum* fed on nine different types of flour (a); the asterisk (*) denotes the internal standard. Peaks labelled 1, 2, and 3 correspond to three major components: 1 – 1-Pentadecene, 2 – cis-9-Tetradecen-1-ol, and 3 – 1-Heptadecene

**FIGURE 2.**
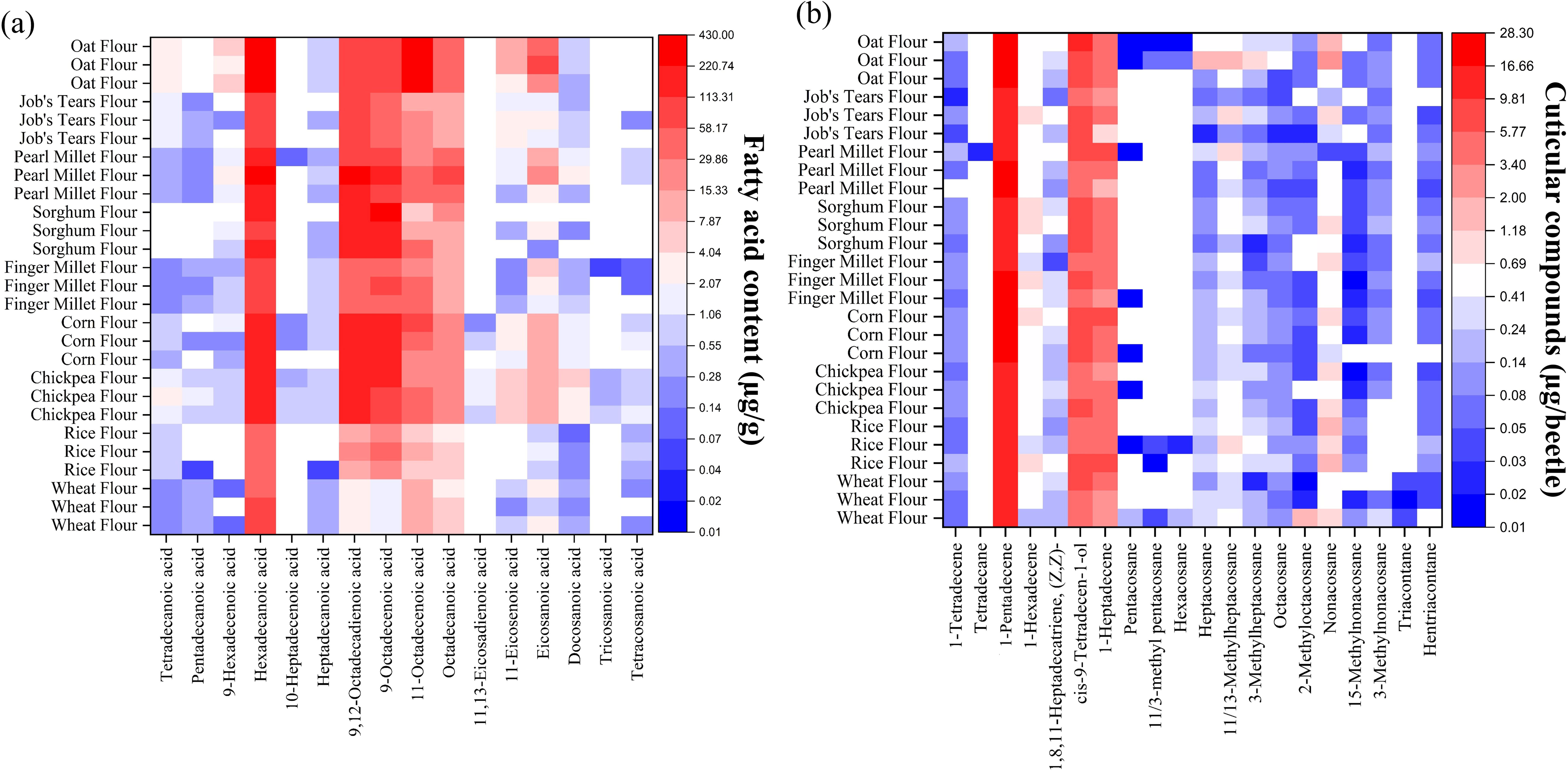
Heatmap showing the quantity of individual fatty acids (µg/g dry weight) across nine different flour types (a). Heatmap representing the quantities of cuticular compounds in different *T. castaneum* populations reared on nine different flours (b). Quantities of each fatty acids/cuticular compounds are represented in Z axis; red (high) to blue (low).

**Table 1.** List of identified fatty acids from nine flour types (n=3).

| Compounds | Rt | Wheat<br>Flour | Rice<br>Flour | Chickpea<br>Flour | Finger<br>Millet<br>Flour | Sorghum<br>Flour | Oat Flour | Corn Flour | Pearl Millet<br>Flour | Job's Tears<br>Flour | p-<br>valu<br>e |
| --- | --- | --- | --- | --- | --- | --- | --- | --- | --- | --- | --- |
| Tetradecanoic<br>acid | 13.1<br>8 | 0.23±0.03 | 0.72±0.1<br>1 | 2.13±0.35 | 0.26±0.01 | 0.00±0.00 | 2.44±0.11 | 0.52±0.11 | 0.40±0.08 | 1.28±0.18 | 0.00<br>1 |
| Pentadecanoic<br>acid | 15.1<br>9 | 0.35±0.03 | 0.02±0.0<br>4 | 1.01±0.21 | 0.31±0.04 | 0.00±0.00 | 0.00±0.00 | 0.05±0.08 | 0.18±0.04 | 0.27±0.03 | 0.00<br>1 |
| 9-Hexadecenoic<br>acid | 16.7<br>8 | 0.06±0.05 | 0.00±0.0<br>0 | 0.79±0.13 | 0.57±0.03 | 0.72±0.63 | 5.00±0.88 | 0.60±0.49 | 2.52±0.81 | 0.08±0.13 | 0.00<br>6 |
| Hexadecanoic<br>acid | 17.1<br>0 | 66.02±12.<br>45 | 42.19±9.<br>77 | 135.08±11.<br>59 | 92.24±10.<br>97 | 124.20±17.<br>00 | 413.03±13.<br>62 | 187.90±21.<br>20 | 206.80±92.<br>47 | 98.42±17.<br>76 | 0.00<br>1 |
| 10-Heptadecenoic<br>acid | 18.5<br>9 | 0.00±0.00 | 0.00±0.0<br>0 | 0.58±0.19 | 0.00±0.00 | 0.00±0.00 | 0.00±0.00 | 0.14±0.12 | 0.04±0.06 | 0.00±0.00 | 0.00<br>8 |
| Heptadecanoic<br>acid | 19.0<br>2 | 0.43±0.05 | 0.02±0.0<br>3 | 0.88±0.14 | 0.59±0.02 | 0.25±0.22 | 0.71±0.09 | 0.49±0.42 | 0.59±0.21 | 0.11±0.19 | 0.01<br>2 |
| Linoleic acid | 20.0<br>3 | 2.61±0.33 | 13.11±3.<br>49 | 177.67±10.<br>60 | 39.12±12.<br>08 | 160.24±15.<br>64 | 67.64±6.42 | 170.16±13.<br>61 | 158.09±77.<br>35 | 80.04±12.<br>71 | 0.00<br>2 |
| Oleic acid | 20.3<br>8 | 1.85±0.19 | 23.66±6.<br>43 | 113.08±19.<br>63 | 56.56±8.6<br>9 | 184.43±63.<br>66 | 101.97±8.5<br>1 | 155.59±18.<br>30 | 133.72±68.<br>74 | 37.24±4.3<br>0 | 0.00<br>2 |
| Elaidic acid | 20.4<br>9 | 9.41±0.64 | 7.93±1.0<br>7 | 42.80±15.9<br>5 | 31.62±1.7<br>2 | 18.38±20.1<br>8 | 238.91±24.<br>75 | 43.70±21.0<br>4 | 39.04±15.5<br>8 | 15.59±0.6<br>4 | 0.01<br>4 |
| Octadecanoic<br>acid | 20.8<br>3 | 4.14±0.21 | 4.36±1.4<br>0 | 24.29±0.98 | 9.54±0.37 | 15.87±3.93 | 49.01±5.74 | 25.66±3.94 | 63.42±30.4<br>3 | 9.20±0.76 | 0.00<br>1 |
| 11,13-Eicosadienoic<br>acid | 23.6<br>4 | 0.00±0.00 | 0.00±0.0<br>0 | 1.45±0.43 | 0.00±0.00 | 0.00±0.00 | 0.00±0.00 | 0.32±0.42 | 0.00±0.00 | 0.00±0.00 | 0.00<br>3 |
| 11-Eicosenoic<br>acid | 23.9<br>9 | 0.62±0.31 | 0.00±0.0<br>0 | 6.29±0.45 | 0.27±0.08 | 0.14±0.23 | 10.22±6.58 | 2.26±0.72 | 1.15±0.77 | 1.97±0.56 | 0.00<br>2 |
| Eicosanoic<br>acid | 24.3<br>1 | 2.85±1.62 | 0.80±0.2<br>8 | 12.52±1.51 | 3.76±1.64 | 0.84±1.27 | 52.07±28.0<br>9 | 9.86±2.06 | 12.69±8.23 | 1.54±0.56 | 0.00<br>4 |
| Docosanoic acid | 27.5<br>3 | 0.36±0.10 | 0.20±0.07 | 4.06±0.49 | 0.46±0.07 | 0.08±0.14 | 0.64±0.19 | 1.43±0.14 | 1.56±1.14 | 0.62±0.15 | 0.005 |
| Tricosanoic acid | 29.3<br>9 | 0.00±0.00 | 0.00±0.00 | 0.50±0.08 | 0.02±0.04 | 0.00±0.00 | 0.00±0.00 | 0.00±0.00 | 0.00±0.00 | 0.00±0.00 | 0.005 |
| Tetracosanoic acid | 31.5<br>4 | 0.13±0.12 | 0.53±0.20 | 1.05±0.25 | 0.08±0.07 | 0.00±0.00 | 0.00±0.00 | 0.69±0.60 | 0.58±0.52 | 0.06±0.11 | 0.03 |

† The table presents the compound names, retention times (Rt), and quantities of the respective compounds. P<0.05, represents statistically significant difference.

Long-chain fatty acids (C > 20), such as docosanoic acid, tricosanoic acid, and tetracosanoic acid, were detected in only a few of the analyzed flour samples, and in very low concentrations. To assess the compositional dissimilarity among the flour samples, a SIMPER (Similarity Percentage) analysis based on Bray-Curtis dissimilarity was performed. The overall Bray-Curtis dissimilarity among the flours was found to be 47.72 (Figure 3b; Supplementary material). The analysis of similarity percentages suggests that each of the nine flour samples possesses a distinct cumulative fatty acid composition (Figure 3d; Supplementary material). The fatty acids contributing most to this dissimilarity were hexadecanoic acid (13.32), 9,12-octadecadienoic acid (12.16), 9-octadecenoic acid (10.79), 11-octadecenoic acid (6.19), octadecanoic acid (2.65), and eicosanoic acid (1.51) (Supplementary material).

**FIGURE 3.**
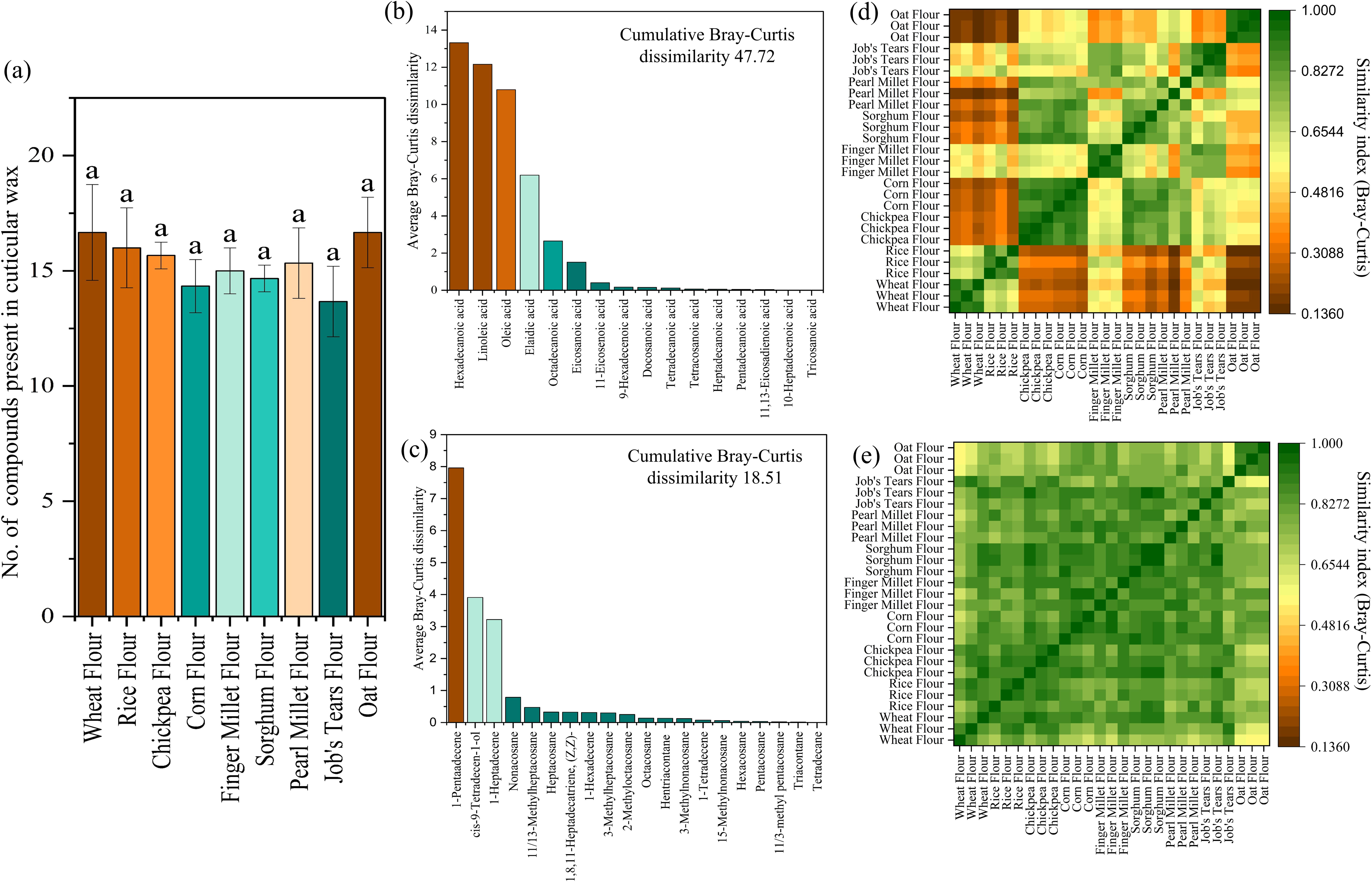
Mean richness of cuticular compounds in different *T. castaneum* populations across diets (a), SIMPER analysis showing average Bray-Curtis dissimilarity contributions of major fatty acids to inter-diet variation (b) and key cuticular compounds of *Tribolium castaneum* fed on different diet groups (c). Heatmap representing the pairwise similarity (Bray-Curtis index) among dietary flours based on their fatty acid profiles (d), and between *T. castaneum* populations reared on different diets based on their cuticular wax profiles (e). Similarity indices are represented in Z axis.

### Chemical characterization of cuticular wax

A total of twenty-five compounds were detected in the cuticular wax of *Tribolium castaneum* (Figure 1b; Table 2). Among these, nonadecane, eicosane, docosane, tricosane, and tetracosane were found only in a single wheat flour sample and were therefore excluded from further analysis. Across all tested samples, 1-pentadecene was the most dominant compound (ranging from 10.74 to 28.22 µg/insect), regardless of the flour source provided (Table 2). This was followed by cis-9-tetradecen-1-ol (3.53– 9.51 µg/insect) and 1-heptadecene (0.72–9.01 µg/insect), which were the second and third most abundant cuticular compounds, respectively (Figure 2b; Table 2). Each analysis typically revealed between 12 and 19 cuticular compounds, with no qualitative variation observed in the compound profiles of *T. castaneum* reared on different flour types (Figure 3a). However, quantitative differences in compound abundance can be observed, though not as significant as fatty acid composition of flours, as supported by the Bray-Curtis similarity matrix (Figure 3e; Supplementary material). To assess these differences in detail, SIMPER analysis was performed. The results indicated a cumulative Bray-Curtis dissimilarity of 18.51 among *T. castaneum* groups reared on different flour sources (Figure 3c; Supplementary material). The compounds contributing most to this dissimilarity were 1-pentadecene (7.96), cis-9-tetradecen-1-ol (3.91), and 1-heptadecene (3.22) (Figure 3c; Supplementary material). Notably, the quantity of 1-pentadecene was significantly higher (H=22.31, p=0.0043) in *T. castaneum* reared on oat flour (26.80 ± 1.41 µg) compared to those grown on wheat (12.13 ± 1.70 µg) and rice flour (12.80 ± 1.04 µg) (Table 2). Although the differences in cis-9-tetradecen-1-ol and 1-heptadecene levels among the different diet groups were not statistically significant, noticeable variation was still observed.

**Table 2.**
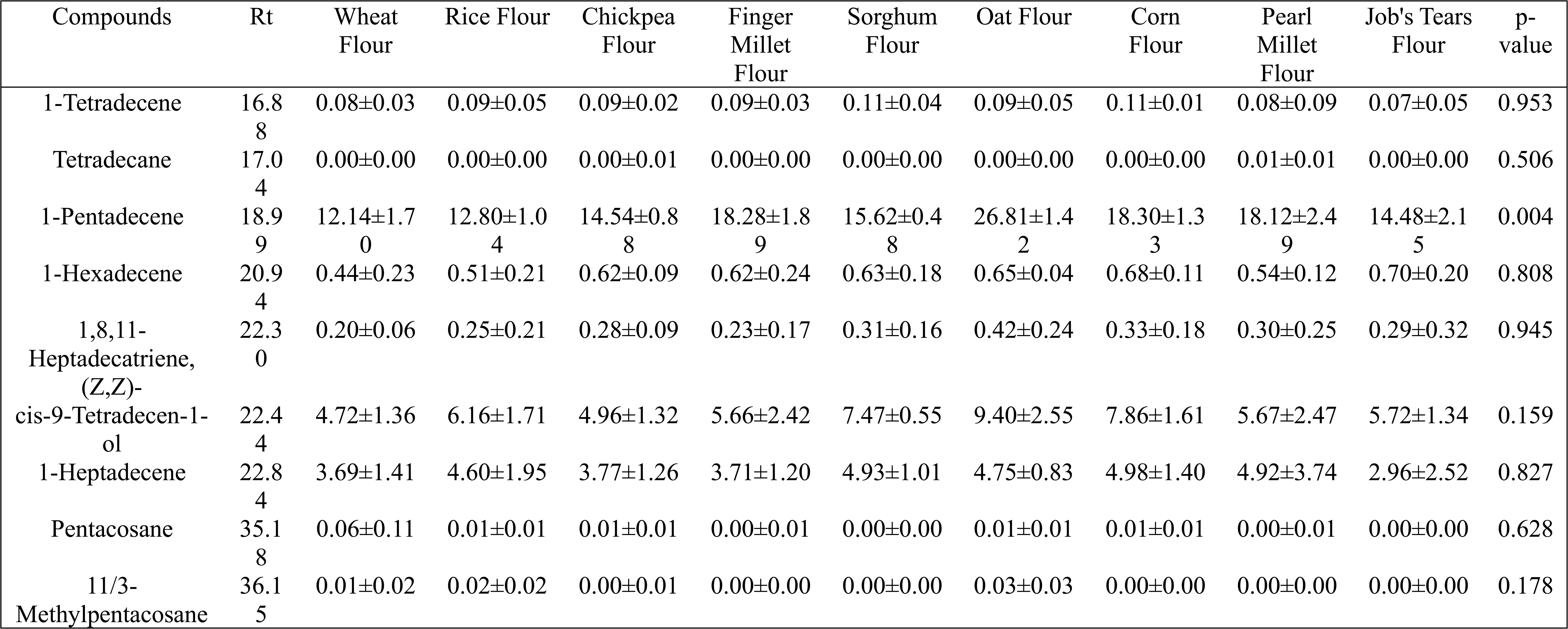
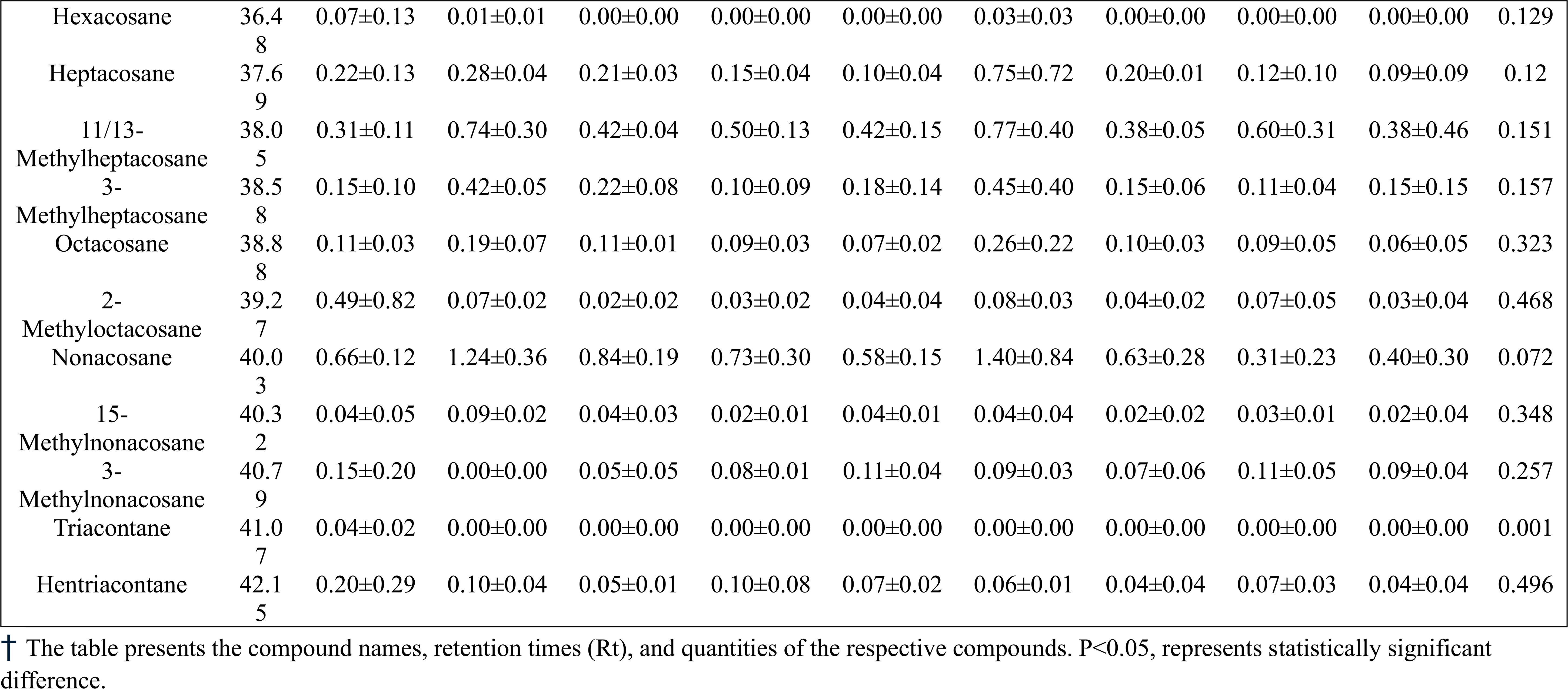
List of identified cuticular wax compounds of *Tribolium castaneum* fed on nine different types of flour (n=3).

### Multivariate Ordination Analysis

Non-metric multidimensional scaling (NMDS) based on the fatty acid profiles of the flour samples was conducted to visualize the clustering pattern among the different flours (Figure 4a; Supplementary material). The analysis revealed that all flour types formed distinct clusters with no prominent overlap, indicating clear differences in their fatty acid compositions. Rice flour, wheat flour, Job’s tears flour, and finger millet flour were localized on the negative axis of NMDS dimension 1, whereas sorghum flour, corn flour, chickpea flour, oat flour, and finger millet flour were positioned on the positive axis of the same dimension. Wheat flour and rice flour were clustered at the extreme negative end of NMDS dimension 1, whereas oat flour was positioned at the extreme positive end (Figure 4a; Supplementary material).

**FIGURE 4.**
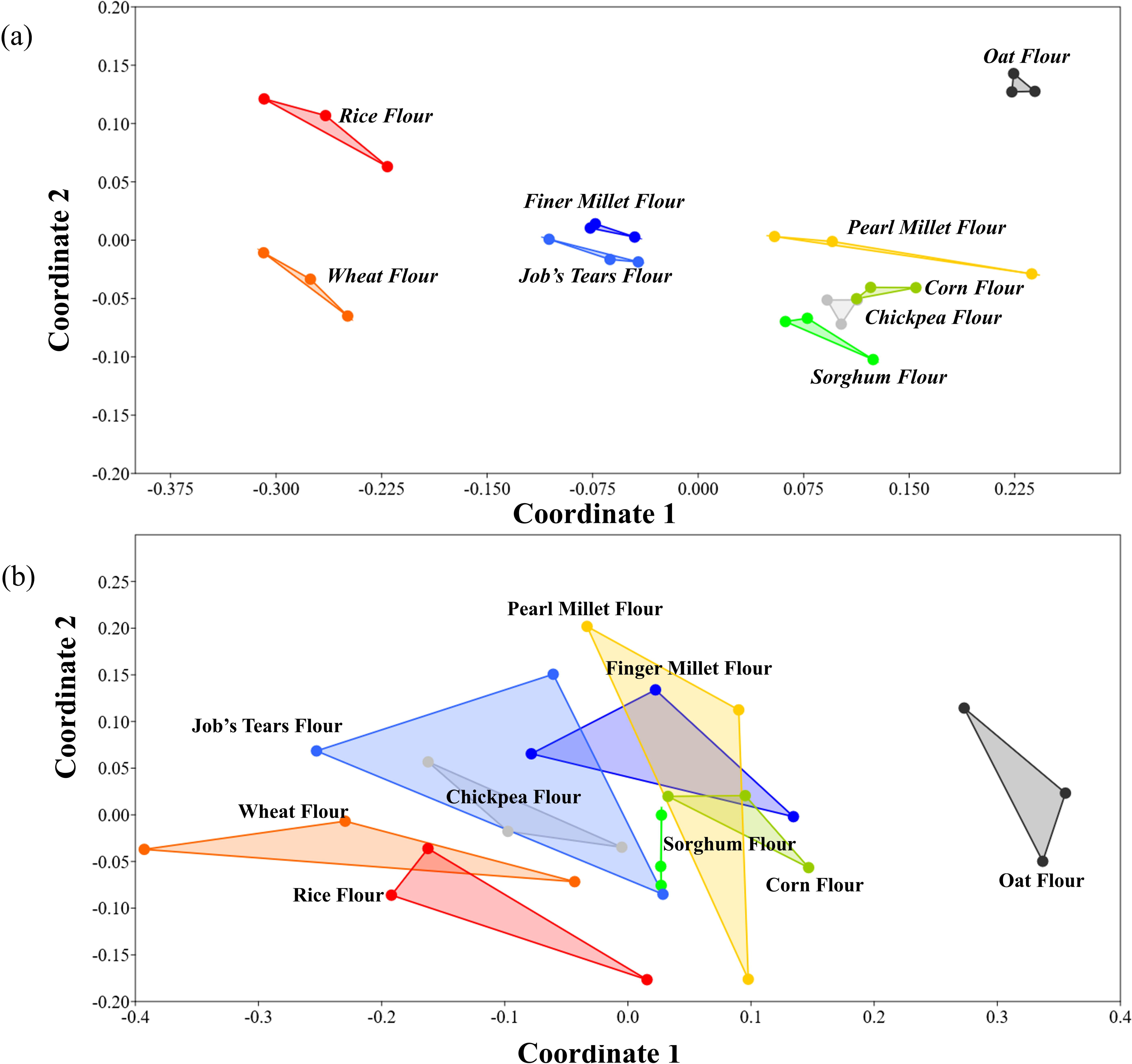
Score plots from non-metric multidimensional scaling (NMDS) analysis, showing separation of dietary flours based on their fatty acid profiles (a), and separation of *T. castaneum* populations reared on different diets based on their cuticular wax profiles (b).

Similarly, to investigate the influence of cuticular chemical composition on the segregation of *T. castaneum* populations reared on the nine aforementioned flours, NMDS was also performed using the chemical profiles of cuticular waxes (Figure 4b; Supplementary material). In contrast to the fatty acid-based clustering of the flours, the *T. castaneum* samples exhibited less distinct segregation, based on cuticular composition, with substantial overlap among individuals reared on different flour types (Figure 4a-b). This suggests a higher degree of intra-group variability in cuticular chemical composition relative to the more discrete differences observed in flour fatty acid profiles. Nonetheless, when evaluating the centroid positions of the clusters, a partially conserved pattern emerged. *T. castaneum* individuals reared on wheat flour, rice flour, Job’s tears flour, and chickpea flour were positioned on the negative axis of NMDS dimension 1, with wheat and rice flour-reared individuals occupying the extreme negative scores. Conversely, individuals reared on sorghum flour, finger millet flour, pearl millet flour, corn flour, and oat flour were clustered on the positive axis, with those associated with oat flour showing the highest positive scores. These findings suggest that, although the segregation was less pronounced, certain broad trends in the cuticular chemical profiles of *T. castaneum* mirrored the clustering pattern observed in the fatty acid profiles of the flours on which they were reared.

### Factor analysis and impact of dietary fatty acids on cuticular chemical profiles of T. castaneum

Another multivariate analysis, i.e., an unsupervised factor analysis or Canonical Analysis Based on Factors (CABFAC) was conducted that relies on the covariance structure among samples, rather than on Bray-Curtis dissimilarity, to further investigate whether clustering patterns of flour samples based on their fatty acid profiles align with the cuticular chemical composition of *T. castaneum* reared on different dietary flour sources. Initially, distinct clusters were observed among the flour samples, similar to those seen in the NMDS analysis. In the factor analysis, 9,12-octadecadienoic acid, 9-octadecenoic acid, and hexadecanoic acid exhibited the highest loadings on Factor 1, while hexadecanoic acid, 11-octadecenoic acid, and 9,12-octadecadienoic acid had the highest loadings on Factor 2—indicating their strong contribution to the variance explained by these latent factors.

In contrast, when the cuticular chemical composition of *T. castaneum* was subjected to factor analysis, although three chemical compounds—1-pentadecene, 1-heptadecene, and cis-9-tetradecen-1-ol— showed substantial loadings, the resulting sample clusters were poorly defined and overlapped considerably, suggesting limited discriminatory power in the latent space based solely on cuticular compounds (Figure 5a).

**FIGURE 5.**
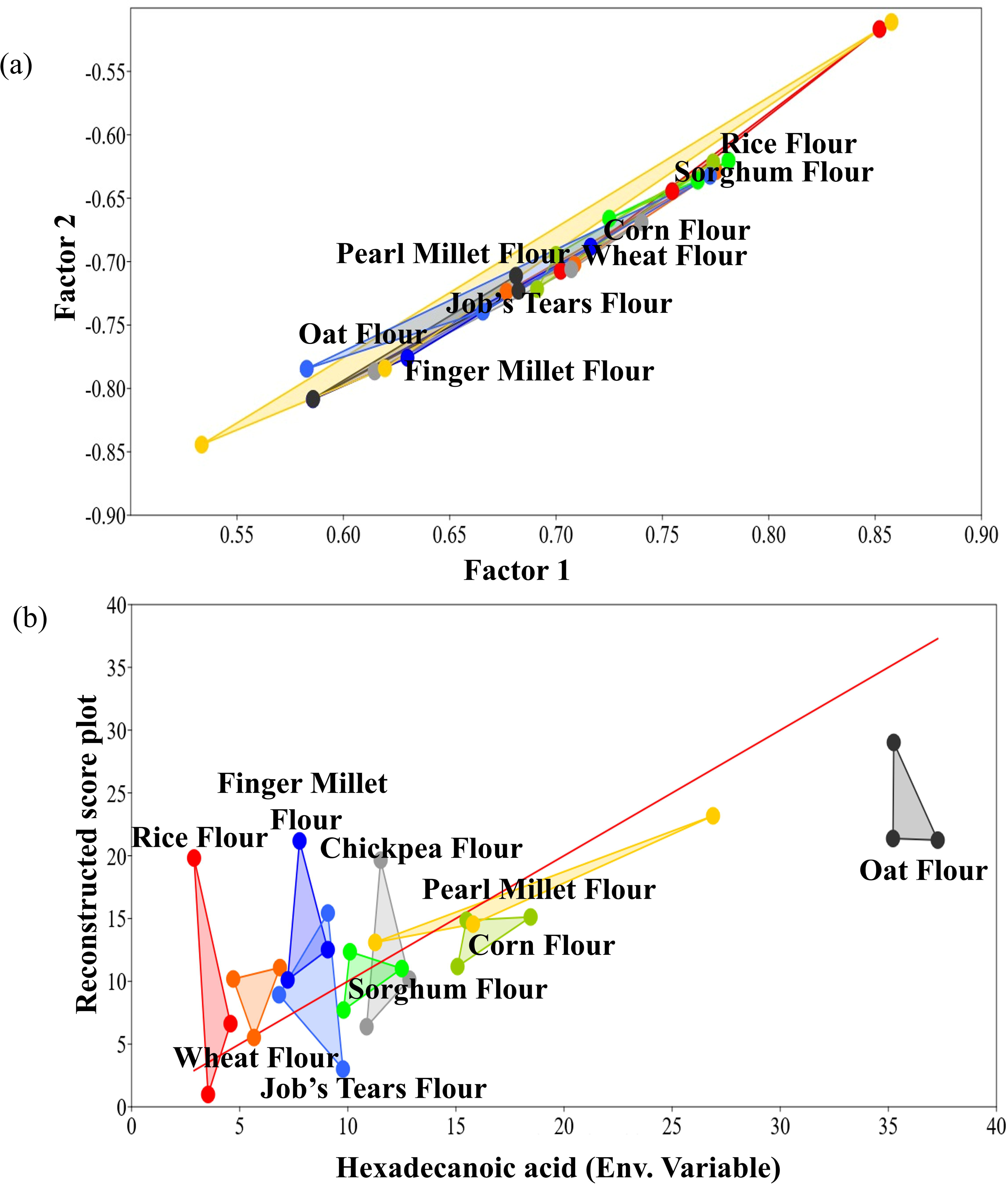
Multivariate analysis correlating dietary fatty acids with cuticular wax composition. Score plot from factor analysis showing the positions of *T. castaneum* populations reared on different flours based on their cuticular wax profiles (a). Reconstructed score plot incorporating dietary fatty acids (Hexadecanoic acid as representative) as environmental variables, illustrating their influence on the clustering of *T. castaneum* populations (b).

To explore this further, individual fatty acid contents in the studied flours were incorporated as exogenous (environmental) variables in a subsequent factor analysis of *T. castaneum*’s cuticular chemical profiles. This integrative approach revealed that only a subset of fatty acids contributed meaningfully to the separation of *T. castaneum* samples based on dietary flour source. Among these, hexadecanoic acid emerged as the most influential variable (Figure 5b), with a normalized RMSE of 4.53% (Supplementary material). These findings suggest that, while not all fatty acids exert equal influence, the concentration of specific fatty acids, particularly hexadecanoic acid, can significantly shape the cuticular chemical composition of *T. castaneum*.

### Multiple correlation analysis

Multiple correlation analysis was conducted to examine how the quantity of individual fatty acids presents in dietary sources (i.e., the provided flours) influences the levels of specific chemical compounds in the cuticular wax of *T. castaneum*. The analysis revealed that not all, but certain chemical constituents of the cuticular wax are significantly correlated with dietary fatty acid content (Figure 6a). For instance, the quantities of 1-pentadecene, cis-9-tetradecen-1-ol, methyl-branched pentacosane, heptacosane, methyl-branched heptacosane, octacosane, and nonacosane in the cuticular wax showed strong positive correlations with specific dietary fatty acids, as illustrated in Figure 6a. Notably, the quantity of 1-pentadecene was strongly and positively correlated with hexadecanoic acid (r=0.90), 9-hexadecenoic acid (r=0.87) (Figure 6b), octadecanoic acid (r=0.64), 11-octadecenoic acid (r=0.87), eicosanoic acid (r=0.71), and 11-eicosenoic acid (r=0.55), and showed a moderate positive correlation with tetradecanoic acid (r=0.45) and heptadecanoic acid (r=0.47) (Supplementary material). A similar, though slightly weaker, correlation pattern was observed for cis-9-tetradecen-1-ol (Figure 6c). In contrast, other wax constituents—namely methyl-branched pentacosane, methyl-branched heptacosane, heptacosane, octacosane, and nonacosane, exhibited strong positive correlations predominantly with eicosanoic acid (r=0.53-0.69) and 11-eicosenoic (r=0.47-0.70), while moderate correlations were also noted with 11-octadecenoic acid (r=0.30-0.50) and tetradecanoic acid (r=0.23-0.39) (Supplementary material).

**FIGURE 6.**
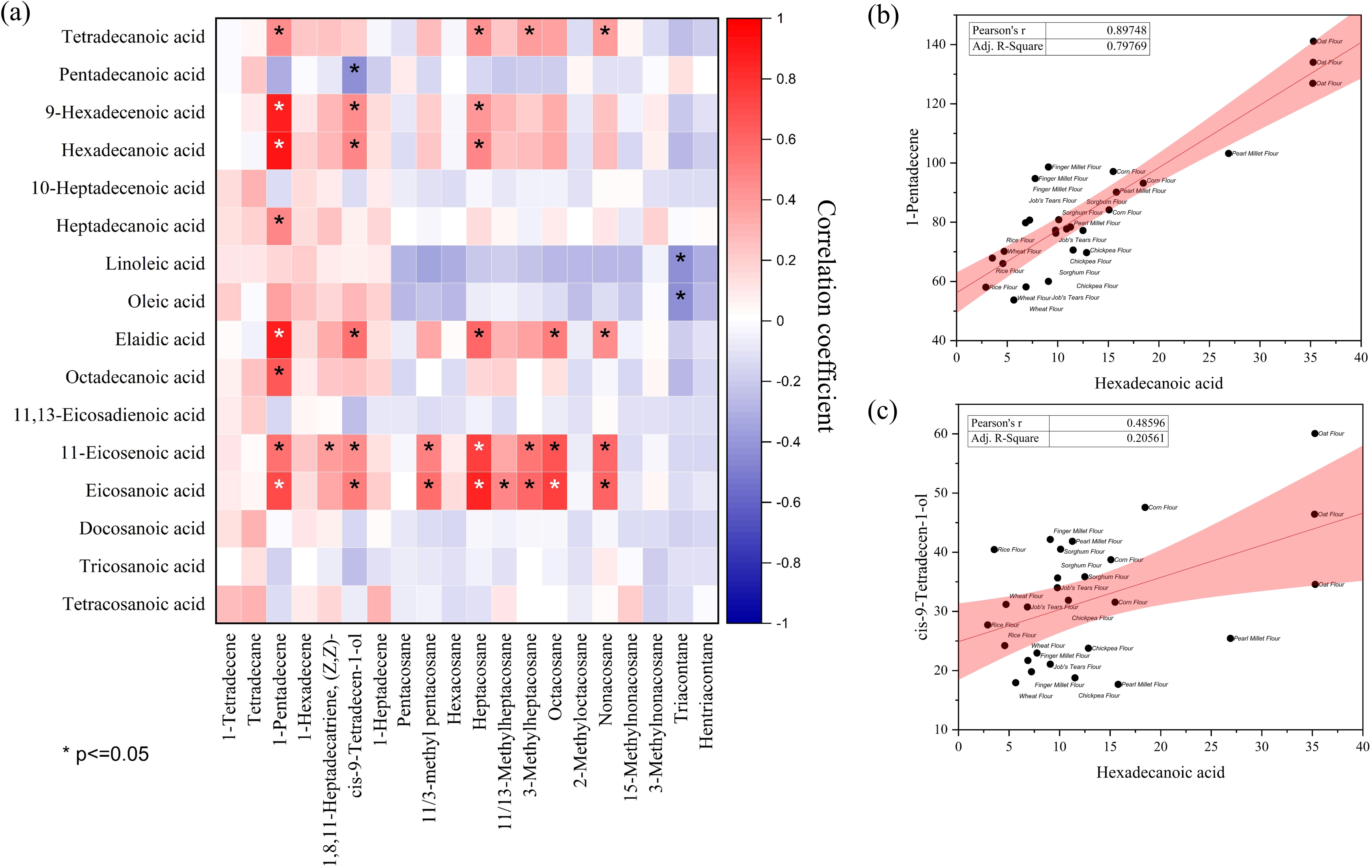
Correlation matrix showing Pearson’s correlation coefficients (r) between dietary fatty acids (rows) and cuticular hydrocarbons in *T. castaneum* populations (columns). Asterisks (*) indicate statistically significant correlations (*p* ≤ 0.05) (a). Scatter plots highlighting the correlations between dietary hexadecanoic acid and the two most abundant cuticular wax components, i.e., 1-pentadecene (b) and *cis*-9-tetradecen-1-ol (c).

## Discussion

In line with previous reports, the present study also identifies 1-pentadecene, cis-9-tetradecen-1-ol, and 1-heptadecene as the dominant compounds in the cuticular wax of *T. castaneum* (Das et al., 2024). Importantly, no substantial qualitative differences in hydrocarbon profiles were observed across populations raised on different diets, corroborating the findings of Otte et al. (2015). This supports the view that the biosynthesis of specific hydrocarbons in insects is primarily governed by their endogenous genetic and metabolic frameworks (Holze et al., 2021). However, quantities of these compounds were found to vary significantly when *T. castaneum* individuals were reared on different dietary sources.

Although previous studies suggest that the composition of cuticular hydrocarbons in the species, including 1-pentadecene, can be significantly influenced by population density, but this variable was controlled in the present study by maintaining a constant population size of 30 individuals through regular flour replacement. This strategy minimized the potential effects of density-dependent variation in hydrocarbon composition. Thus, the given diet was the only external factor varied in this study, and it can be inferred that dietary components, especially dietary fatty acid, may still influence relative abundances of CHCs in cuticular wax of *T. castaneum* to some extent.

Several studies (Buczkowski et al., 2005; Buczkowski and Silverman, 2005; Fedina et al., 2012; Liang and Silverman, 2000; Otte et al., 2015; Pennanec’h et al., 1997; van Wilgenburg et al., 2022b) have documented associations between diet and cuticular hydrocarbons, although most do not specify links between individual macromolecules and hydrocarbon profiles. Exceptions include the works of Pennanec’h et al. (1997) and Otte et al. (2015), which suggest a direct relationship between dietary fatty acids and CHC biosynthesis. For instance, Otte et al. (2015) demonstrated in *Phaedon cochleariae* (mustard leaf beetle) that the ratio of mono- to di-unsaturated CHC in cuticular wax is correlated with the corresponding ratio of mono- to di-unsaturated fatty acids in their host plant. Likewise, Pennanec’h et al. (1997) showed that *D. melanogaster* fed on radiolabelled fatty acids exhibited elevated levels of radiolabel incorporation into specific hydrocarbons, i.e., 7-tricosene (C23) in males and 7,11-heptacosadiene (C27) in females. Their findings suggested that dietary even-chain fatty acids, such as Tetradecanoic (C14) and hexadecanoic acids (C16) (mainly C14), are precursors to aforesaid odd-chain hydrocarbons via initial decarboxylation followed by chain elongation (Pennanec’h et al., 1997).

In our study, SIMPER analysis revealed a Bray–Curtis dissimilarity of 47.72 among the nine flour samples based on their fatty acid profiles, while the same analysis yielded a Bray–Curtis distance of 18.51 among *T. castaneum* populations reared on these flours. This suggests that although dietary fatty acids are not the sole determinants, but they contribute ascertainably to the variation in cuticular hydrocarbon composition. NMDS analysis further supports this conclusion, showing a weak but consistent clustering pattern in case of flours (based on fatty acid profiles) and the corresponding *T. castaneum* populations (based on CHC profile). Factor analysis highlighted the role of specific dietary fatty acid’s concentrations (as environmental variables) that facilitate clustering of *T. castaneum* populations based on CHC (cuticular hydrocarbon) profiles. Both factor analysis and subsequently, Pearson’s correlation revealed a strong positive correlation between dietary hexadecanoic acid (C16) and the abundance of 1-pentadecene (C15) in the insect cuticle. Previous studies (Blomquist et al., 1987; Pennanec’h et al., 1997) suggest that such transformation could occur via one-step decarboxylation to yield n-pentadecane, which may subsequently undergo terminal desaturation. Alternatively, the transformation of hexadecanoic acid (C16) directly to 1-pentadecene (C15) could involve cytochrome P450 fatty acid decarboxylases homolog genes in *T. castaneum*, similar to those identified in *Jeotgalicoccus* sp., which can facilitate terminal olefination in n-alkane (Liu et al., 2014; Rude et al., 2011).

In addition to hexadecanoic acid (C16), abundance of 1-pentadecene (C15) in *T. castaneum* populations was found to correlate with other even-chain fatty acids, including tetradecanoic (C14), octadecanoic (C18), and eicosanoic acids (C20). Among these, octadecanoic (C18) and eicosanoic acids (C20) may undergo multistep decarboxylation and desaturation, while tetradecanoic acid (C14) may first be elongated before undergoing decarboxylation. The elongation of tetradecanoic acid (C14) appears to support the biosynthesis of long-chain hydrocarbons such as heptacosane (C27), branched heptacosane (brC27), and Nonacosane (C29). This is consistent with the findings of Pennanec’h et al. (1997), who observed that tetradecanoic acid (C14) is more efficiently incorporated into long-chain cuticular hydrocarbons like 7-tricosene (C23) and 7,11-heptacosadiene (C27) in *D. melanogaster* than are hexadecanoic acid (C16) or octadecanoic acid (C18). Similarly, strong correlation between long-chain dietary fatty acids, such as eicosanoic acid (C20), and long chain CHCs in cuticular wax of *T. castaneum*, alongside the lack of such correlation of long-chain fatty acids with short-chain CHCs further indicated that long-chain fatty acids primarily serve as precursors for long-chain hydrocarbon biosynthesis via elongase-mediated pathways.

## Conclusion

Although this study does not comprehensively elucidate the metabolic pathway connecting dietary fatty acids to specific cuticular hydrocarbons in *T. castaneum*, the findings of this study provide strong empirical support for the hypothesis that, beyond endogenous metabolic regulation, dietary fatty acid composition plays a significant role in modulating the insect’s cuticular hydrocarbon profile.

## Acknowledgements

This research was conducted at the Semiochemicals and Lipid Laboratory, Department of Life Sciences, Presidency University, Kolkata. We extend our heartfelt gratitude to the head of this lab, Prof. Mousumi Poddar Sarkar, for her invaluable guidance and support. We also acknowledge the DBT-BUILDER programme, Government of India (BT/INF/22/SP45088/2022, dated 17.02.2022), for providing the GC-MS facilities at the Department of Life Sciences, Presidency University, Kolkata.

## Data availability statement

The author confirms that the data supporting the findings of the study are available within the article and raw data that support the findings of this study are available from the corresponding author upon request.

## Ethics statement

This study does not require ethical approval.

## Author contributions

**Subhadeep Das**: Investigation, Methodology, Formal analysis, Writing-Original draft preparation, Data curation, Writing-Reviewing and Editing. **Oishika Chatterjee**: Data curation, Reviewing, Methodology. **Riya Saha**: Data curation, Reviewing, Methodology. **Pritom Das**: Methodology, Reviewing. **Avisekh Dolai**: Formal analysis. **Sourav Manna**: Conceptualization, Visualization, Writing, Methodology, Validation.

